# Genome quality variation across Scyphozoa and the comparative distribution of retinoid- and AhR-related gene families

**DOI:** 10.64898/2026.04.22.720242

**Authors:** Yeong-Jun Park, Nayoung Lee, Yejin Jo, Seungshic Yum, Kae Kyoung Kwon

## Abstract

Scyphozoan jellyfish have a complex life cycle that includes a characteristic transition known as strobilation. Retinoid signaling has been suggested to be involved in jellyfish metamorphosis and development. However, the genomic basis of signaling pathways associated with metamorphosis has not been sufficiently compared at the class level. Experimental studies have reported that indole compounds can induce metamorphosis in some jellyfish species. Indole- and tryptophan-derived metabolites are known to function as ligands for the aryl hydrocarbon receptor (AhR) in other organisms. However, the potential role of AhR signaling in jellyfish metamorphosis has not been previously explored. We compared the distribution of retinoid- and AhR-associated gene families across multiple scyphozoan genomes. This analysis aimed to characterize their distribution patterns in relation to signaling pathways associated with development and environmental responses. A standard gene prediction and annotation pipeline was applied to 20 species from 21 publicly available scyphozoan reference genome assemblies retrieved from the NCBI database. The distribution and copy number of these gene families were compared across species. Retinoid-associated gene families were detected across almost all Scyphozoa genomes, and core components of AhR signaling (AhR, ARNT) were identified in most species. These results suggest that scyphozoan genomes contain genetic components of retinoid- and AhR-related signals. This study presents the distribution of gene families related to developmental signaling across Scyphozoa using a comparative genomic approach. It does not imply direct functional involvement of retinoid or AhR signaling, but instead focuses on potential signaling pathways at the genome level. It also provides an overview of currently available scyphozoan genomic data. These findings provide a basis for future hypothesis generation and functional validation in jellyfish metamorphosis research.

## Introduction

Scyphozoa exhibit a complex life cycle that alternates between sessile and pelagic stages, including polyp, strobila, ephyra, and medusa stages [1]. A key developmental transition in this cycle is strobilation, during which the polyp transforms into the ephyra [1]. The sessile polyp and the pelagic ephyra show marked morphological differences. This transition is accompanied by extensive tissue reorganization and changes in physiological state during strobilation [2]. Strobilation is therefore regarded as one of the most important developmental turning points in the scyphozoan life cycle [1]. In addition, scyphozoan metamorphosis generates distinct biological states within the same individual and is sensitive to environmental signals [3]. Therefore, investigating related chemical or environmental cues is important for understanding metamorphosis [3]. Such large-scale developmental changes require sophisticated molecular regulatory systems, and metamorphosis is considered the result of controlled developmental programs [1]. However, molecular evidence for signal recognition and transduction in the host remains limited [4].

To date, most studies have focused on morphological and developmental interpretations [1]. More recently, many studies have relied on observations of phenomena following exposure to chemicals [4]. However, studies at the molecular level, particularly genome-based comparative analyses related to metamorphosis, remain insufficient. Importantly, class-wide comparisons of conserved developmental signaling systems in Scyphozoa remain limited [5]. To achieve this, an overall assessment of genome assemblies and gene annotation status must first be carried out first.

Among the signaling pathways associated with jellyfish metamorphosis, retinoid signaling has been the most extensively studied pathway associated with jellyfish metamorphosis, and a subset of genes has been reported, such as retinoid X receptor (RXR), retinol dehydrogenase 1 (RDH1), and retinol dehydrogenase 2 (RDH2); however, the complete set of genes within this pathway has not yet been fully characterized [1,4]. In addition, microbial β-carotene has been reported to potentially influence metamorphosis [6], suggesting that β-carotene cleavage genes such as BCO1 and BCO2 may be involved in strobilation. Through these genes, the host retinoid pathway may be linked to microbial β-carotene, indicating a potential extension of host-microbe metabolic interactions. Beyond retinoid compounds, indole-derived compounds have also been reported to influence metamorphosis in some jellyfish species; nevertheless, the genetic basis underlying this process has not yet been proposed [4]. In this regard, indole- and tryptophan-derived metabolites have been shown to function as ligands for aryl hydrocarbon receptor (AhR) in other animals [7,8]. Consequently, a potential association between AhR and indole compounds may exist in jellyfish, and AhR may function as a receptor involved in environmental responses. However, the genetic composition and distribution of the AhR signaling system in jellyfish have rarely been discussed.

Although Scyphozoa is estimated to comprise more than 200 species [1], only genome assemblies from 21 species (the majority belonging to two orders inhabiting surface area) are currently available in the NCBI database, and no complete genome assembly has yet been reported [5]. Furthermore, genome assemblies released in the NCBI database may have variable annotation quality and considerable differences in gene completeness due to sequencing quality and analytical methods [5,9]. This situation makes genome-based comparative analyses across species difficult. Nevertheless, class-level comparisons are necessary to understand jellyfish biological characteristics, as they provide a basis for shared features from a genomic perspective.

Genome analyses across diverse species should be conducted prior to studies of the strobilation process, which is closely related to jellyfish blooms [10]. For this purpose, genome assembly quality should be compared, and gene distributions should be evaluated while considering assembly quality [5,9]. In addition, gene prediction and annotation should be performed and compared according to standardized criteria [11,12]. As the current stage, rather than seeking to infer functional conclusions about metamorphosis-related genes through comparative genomic analyses, it is more appropriate to focus on examining whether key candidate genes are broadly present across Scyphozoa and to compare their copy number distributions [13,5].

In this study, we evaluated the assembly quality and gene completeness of available Scyphozoa genome assemblies to assess whether they are sufficient for comparative analyses, and we summarized the current status of genomic resources [9]. In addition, we examined the presence of signaling-related genes associated with metamorphosis, based on biological evidence previously reported across diverse species [13]. Furthermore, the distributions of key candidate genes were compared at the class level to interpret previously scattered findings as shared characteristics across Scyphozoa [5]. Based on these analyses, we aimed to provide a comparative genomic framework for future functional studies of metamorphosis.

## Materials and Methods

### Genome data collection

In January 2026, 21 publicly available Scyphozoa genome assemblies were downloaded from the NCBI GenBank database [14]. The assembly levels were chromosome- or scaffold-level, and the corresponding GenBank accession numbers are provided in Table 1. Genome quality assessment was performed for all species, whereas gene prediction and annotation were conducted for 20 species, excluding one species. Owing to its large genome size (approximately 4.8 Gbp), gene prediction for this species could not be completed because of insufficient memory resources on a public server environment. The species was included in the QUAST- and BUSCO-based quality assessment but excluded from the gene prediction stage.

**Table 1.**
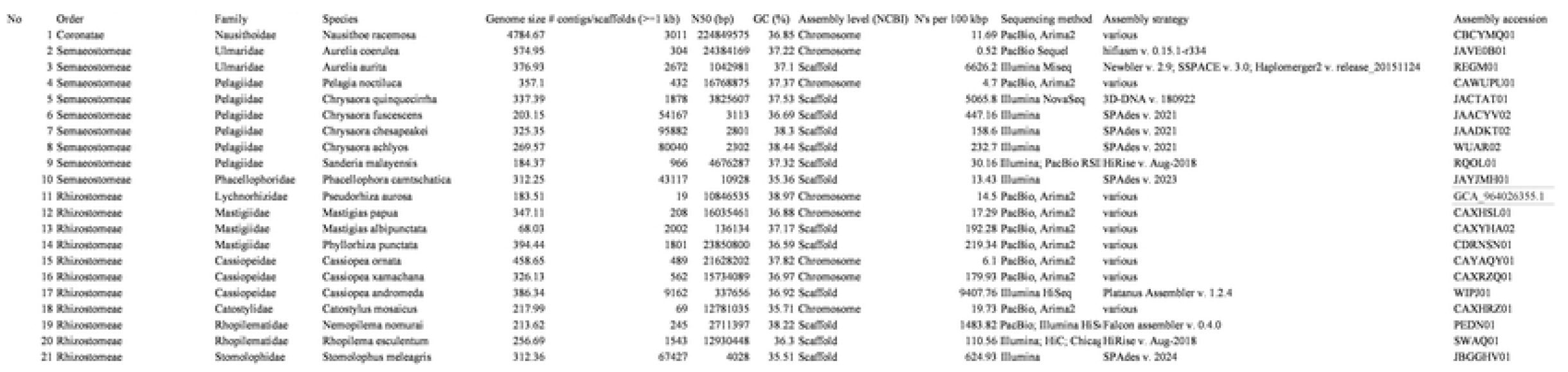
Summary of genome assembly statistics for 21 scyphozoan species.

### Genome quality assessment

To assess the overall assembly quality of all 21 species, QUAST v5.3.0 was used [15]. Gene-based completeness was evaluated using BUSCO v6.0.0 with the metazoa_odb10 lineage dataset, running in genome mode (-m genome) [9]. BUSCO completeness was determined based on the proportion of complete BUSCOs, including single-copy and duplicated categories [9]. Default parameters were used for BUSCO analysis unless otherwise specified.

### Gene prediction and functional annotation

Gene prediction and functional annotation were performed for 20 species. Gene prediction was conducted using funannotate v1.8.17, with the AUGUSTUS species model of *Aurelia aurita*, which represents a comparatively well-annotated scyphozoan genome among the available AUGUSTUS models [16]. Analyses were performed using the default funannotate pipeline. Protein evidence (UniProt/Swiss-Prot; uniprot_sprot.fasta) was incorporated for all species during gene prediction, and no taxonomic filtering was applied at this stage [16]. As output, predicted protein sequences were obtained. Functional annotation was carried out using ortholog-based approaches implemented in eggNOG-mapper v2.1.12 with the eggNOG database v5.0.2 [17]. Genes of interest were identified based on ortholog assignments in the eggNOG-mapper output.

### Identification and comparative analysis of AhR- and retinoic acid-related gene families

The genes summarized in Table S1 and S2 were selected to describe the distribution of target gene families across scyphozoan species. Rather than evaluating the completeness or activity of specific pathways, the present analysis focused on comparing the presence and copy number of orthologous gene families identified across species based on orthology-supported annotations [13].

Table S1 summarizes gene families previously reported to be associated with retinoid metabolism and retinoid-related signaling processes [1]. Genes annotated as RXRA-like were included based on their classification within the corresponding functional category in the eggNOG annotation framework [17]. With respect to carotenoid cleavage, the BCO1 and BCO2 gene families, which have been reported to catalyze the production of retinal or retinol precursors from β-carotene, were selected for comparison [18]. In addition, genes belonging to RDH-like and SDR families, which are involved in retinol-retinal interconversion, were included as representative components of retinoid metabolic pathways [19]. Finally, ALDH1A1 was selected as a representative member of gene families associated with the oxidation of retinal to retinoic acid [20].

Table S2 includes gene families that have been reported to be directly or indirectly associated with the aryl hydrocarbon receptor (AhR) [7]. Core components related to AhR-associated signaling, including AhR, ARNT, and AHRR, have been described across diverse animal taxa and were therefore included for comparative assessment [21].

### Software and parameters

The software used for the analyses included QUAST v5.3.0 [15], BUSCO v6.0.0 [22], funannotate v1.8.17 [16], eggNOG-mapper v2.1.12 [17], and BLASTP (NCBI web-based tool) [23]. The databases used in this study included the BUSCO lineage dataset metazoa_odb10, the eggNOG database v5.0.2 [24], as well as the UniProt/Swiss-Prot and NCBI nr databases [25,26]. BUSCO analyses were conducted in genome mode (-m genome). For gene prediction using funannotate, the default workflow was applied, with *Aurelia aurita* specified as the AUGUSTUS species model [16,27]. Predicted protein sequences generated by funannotate were used as input for eggNOG-mapper for orthology-based functional annotation, and protein-protein homology searches were performed using BLASTP. Unless otherwise specified, default parameters were used for all analyses.

## Results

### Genome assembly quality and completeness of 21 scyphozoan species

The statistical characteristics of the 21 scyphozoan genome assemblies are summarized in Table 1. Genome sizes were distributed as follows: one species had a genome size below 100 Mbp, two species ranged from 100 to 200 Mbp, five species from 200 to 300 Mbp, ten species from 300 to 400 Mbp, and three species exceeded 400 Mbp. Among the analyzed species, *Nausithoe racemosa* exhibited the largest genome assembly, with a total size of approximately 4.8 Gbp.

The number of contigs or scaffolds longer than 1 kb varied widely among species. While *Pseudorhiza aurosa* and *Catostylus mosaicus* contained only a few dozen contigs/scaffolds, several species, including *Chrysaora fuscescens, Stomolophus meleagris, Phacellophora camtschatica, Chrysaora chesapeakei*, and *Chrysaora achlyos*, showed tens of thousands of contigs/scaffolds. N50 values also spanned a broad range, from 2,302 bp to a maximum of 224,849,575 bp. Although sequencing platforms and assembly strategies differed among species, GC content showed a relatively narrow distribution, ranging from 35.36% to 38.97% (Table 1).

BUSCO analysis using the metazoa_odb10 dataset showed differences in the composition of BUSCO categories (S, D, F, and M) among species (Figure 1, Table S3). In many genome assemblies, complete and single-copy BUSCOs (S) accounted for more than 80% of the total. In contrast, some species exhibited relatively high proportions of complete and duplicated BUSCOs (D), with two species showing D values exceeding approximately 20%. In addition, several genomes displayed elevated proportions of fragmented (F) or missing (M) BUSCOs.

**Figure 1.**
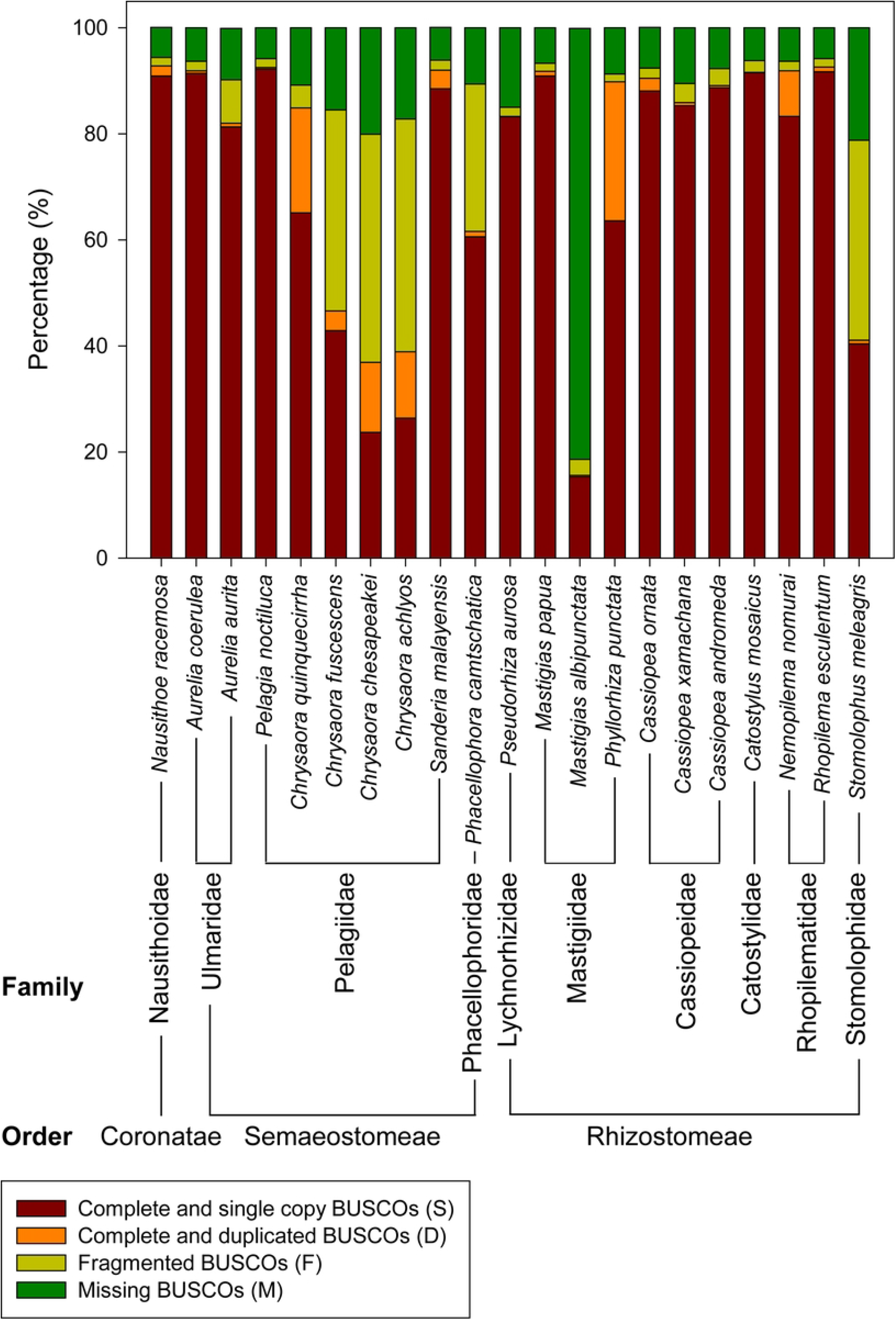
BUSCO completeness of 21 scyphozoan genome assemblies.

Based on complete BUSCOs (C), many of the analyzed species retained a substantial fraction of the metazoan core gene set, whereas a subset of genomes showed C values in the ∼60% range or lower (Table S3).

### Gene prediction and annotation summary across species

Based on gene prediction results generated using funannotate, the number of predicted protein-coding genes across the analyzed scyphozoan genomes ranged from approximately 16,000 to 42,000, with noticeable variation among species (Table 2). Among these, *Mastigias albipunctata* showed a markedly lower number of predicted genes compared to other species. Predicted gene numbers were estimated by counting the number of protein sequences generated in the final funannotate protein output files.

**Table 2.** Gene prediction and annotation summary across 20 scyphozoan.

| No. | Species | Predicted genes | Mean gene length (bp) | Mean exons per gene | eggNOG annotation coverage (%) |
| --- | --- | --- | --- | --- | --- |
| 1 | *Aurelia coerulea* | 20676 | 5620.72 | 5.31719 | 54.7785 |
| 2 | *Aurelia aurita* | 16460 | 6701.34 | 6.70208 | 62.5091 |
| 3 | *Pelagia noctiluca* | 28196 | 2281.26 | 3.49576 | 56.0186 |
| 4 | *Chrysaora quinquecirrha* | 23753 | 4738.46 | 5.87796 | 61.8617 |
| 5 | *Chrysaora fuscescens* | 34790 | 1559.59 | 3.28246 | 59.1865 |
| 6 | *Chrysaora chesapeakei* | 42425 | 1444.9 | 2.98319 | 57.3765 |
| 7 | *Chrysaora achlyos* | 26594 | 1615.42 | 3.34992 | 62.0365 |
| 8 | *Sanderia malayensis* | 24006 | 3231.6 | 6.82595 | 59.3477 |
| 9 | *Phacellophora camtschatica* | 42101 | 2614.47 | 3.93861 | 59.2884 |
| 10 | *Pseudorhiza aurosa* | 16883 | 5550.2 | 6.83431 | 64.9825 |
| 11 | *Mastigias papua* | 27393 | 5011.91 | 7.29537 | 55.1163 |
| 12 | *Mastigias albipunctata* | 2771 | 7285.44 | 8.01512 | 61.4219 |
| 13 | *Phyllorhiza punctata* | 32760 | 5252.68 | 7.40355 | 51.6911 |
| 14 | *Cassiopea ornata* | 33433 | 4826.47 | 5.9242 | 49.9357 |
| 15 | *Cassiopea xamachana* | 27980 | 7199.92 | 6.562 | 48.6133 |
| 16 | *Cassiopea andromeda* | 32015 | 5247.88 | 6.01276 | 53.0564 |
| 17 | *Catostylus mosaicus* | 20302 | 5599.59 | 8.20313 | 63.9494 |
| 18 | *Nemopilema nomurai* | 22004 | 5204.85 | 7.93533 | 62.3659 |
| 19 | *Rhopilema esculentum* | 23220 | 7838.94 | 7.38137 | 53.8286 |
| 20 | *Stomolophus meleagris* | 35686 | 2080.38 | 3.34036 | 50.6025 |

The mean gene length varied across species, ranging from approximately 1,400 bp to 7,800 bp. Mean gene lengths were calculated based on gene feature coordinates in the funannotate-generated GFF3 files. In addition, the mean number of exons per gene ranged from approximately 3 to 8, indicating differences in the structural characteristics of predicted gene models among species.

Functional annotation using eggNOG-mapper was performed on protein-coding genes, excluding tRNA, rRNA, and pseudogenes. A proportion of predicted proteins was assigned orthology-based functional annotations, and annotation coverage varied across species, ranging from approximately 49% to 65% (Table 2).

### Distribution of selected retinoid- and AhR-associated gene families across scyphozoan genomes

Based on gene prediction and annotation results, we examined the species-specific distribution of selected retinoid- and AhR-associated gene families across 20 scyphozoan genome assemblies. One species exhibiting very low BUSCO gene completeness was excluded from further analysis due to limited reliability in target gene identification, and gene distribution patterns were compared among the remaining 19 species (Table S3).

With respect to retinoid signaling, the retinoic acid receptor (*RXR*) was identified in most scyphozoan genomes, except for two species. Genes involved in β-carotene degradation, including *BCO1* and/or *BCO2*, were detected in all analyzed species. In addition, *RDH-like* and *SDR16C5* genes, which are thought to be involved in the conversion of carotenoid-derived precursors into retinol or retinal, were widely distributed across species. The distribution of *RLBP1*, a retinal-binding protein, and *ALDH1A1*, which catalyzes the oxidation of retinal to retinoic acid, further indicated that gene families related to retinal-to-retinoic acid conversion are present in many scyphozoan genomes (Figure 2A).

**Figure 2.**
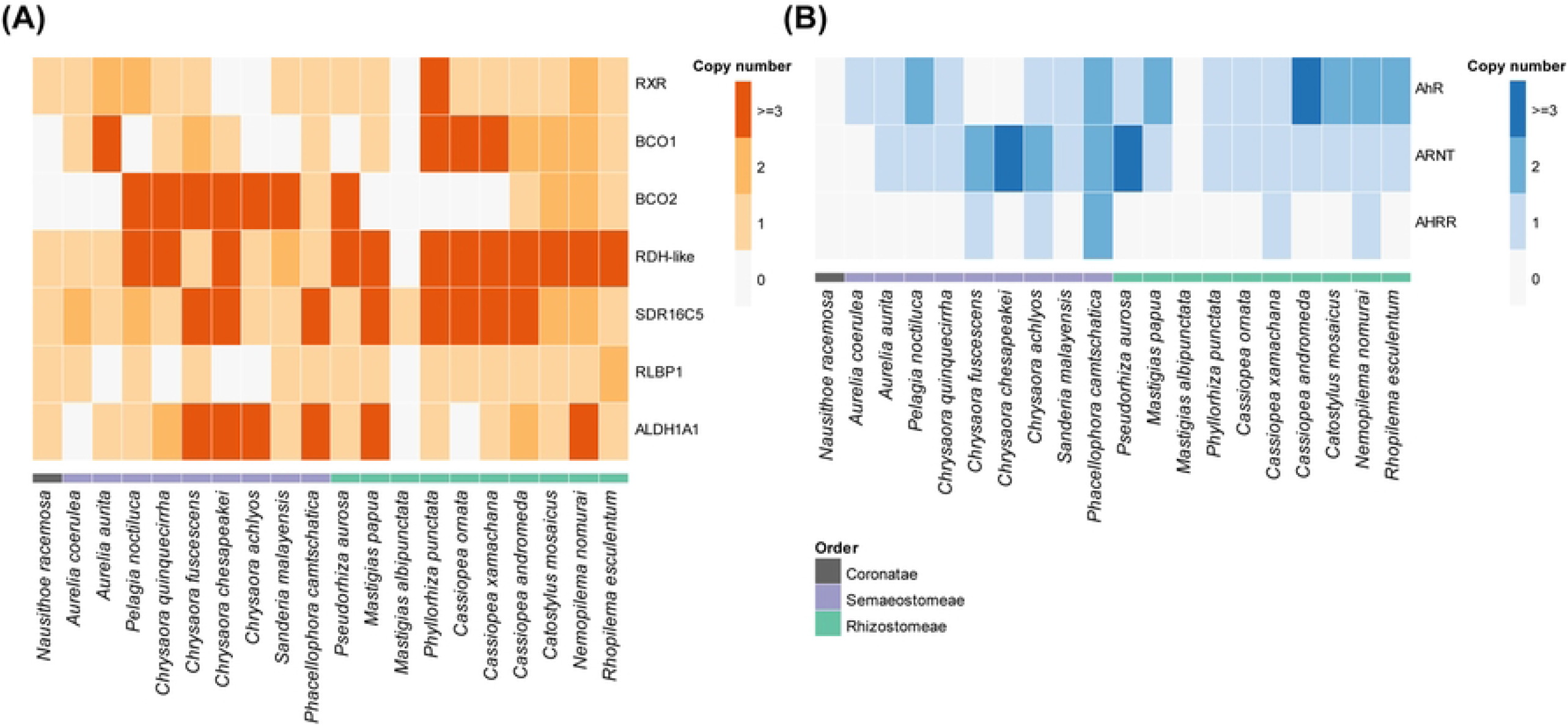
Distribution of conserved retinoid- and AhR-associated gene families across scyphozoan genomes. (A) Heatmap showing the copy number distribution of retinoid signaling-related gene families, including RXR, BCO1, BCO2, RDH-like, SDR16C5, RLBP1, and ALDH1A1, across 20 scyphozoan genomes. (B) Heatmap showing the copy number distribution of core components of AhR signaling pathway, including AhR, ARNT, and AHRR.

Analysis of AhR-related gene families showed that the aryl hydrocarbon receptor (*AhR*) was detected in most genomes, with the exception of two species. The aryl hydrocarbon receptor translocator (*ARNT*), which forms a functional heterodimer with AhR and participates in transcriptional regulation, was also present in all but one genome. In contrast, *AHRR*, a known negative regulator of AhR signaling in vertebrates, was absent from most scyphozoan genomes (Figure 2B).

## Discussion

Scyphozoa reference genome assemblies vary in quality and completeness across species [5]. This variation can limit the interpretation of genome-based comparative analyses at the class level [9]. In practice, BUSCO completeness, assembly contiguity, and gene model complexity varied widely among species. To enable reliable comparisons of gene presence/absence and relative distributions, the same annotation pipeline was applied across all genomes, with additional options for improving the reliability of gene prediction and annotation as described in the Methods [11]. Some observed gene absences may reflect not only true biological absence but also assembly fragmentation or annotation errors [9].

No clear presence-absence patterns at the order level were observed for the retinoid- and AhR-related gene families examined in this study. These genes were detected across most species, indicating that they are widely distributed Scyphozoa rather than lineage-specific features.

This pattern was consistently observed in retinoid signaling-related gene families. Genes involved in the production of 9-cis-retinoic acid, a known ligand of the retinoid X receptor (RXR), including *BCO, RDH-like, ALDH1A1-like* genes, together with *RXR*, were selected as components of retinoid signaling, and their genomic presence was examined [18,19,20]. Copy number variation was detected for these genes, whereas core gene sets were consistently detected across species. Retinoid-associated gene repertoires are not species-specific but appear to represent shared genomic features across Scyphozoa.

AhR signaling core components, AhR and ARNT, were detected in most Scyphozoa genomes. The presence of these genes suggests a genomic potential for responding to external chemical stimuli [7]. Potential sources of AhR ligands may include endogenous synthesis or microbial indole compounds [7,8]. However, these possibilities were not examined in this study. The uneven distribution of AHRR suggests that its well-known role as a negative regulator of the pathway may differ in Scyphozoa. This pattern may be partly explained by the absence of AHRR in several species, although the functional implications remain unclear.

The presence of genes associated with the two signaling systems described above suggests a potential genomic framework that may support developmental signaling at the genomic level. Although genome-based analyses indicate potential involvement of these genes, such analyses do not provide information on the timing, location, or mechanisms of gene activation [28]. Stage-specific activation is expected to depend more on transcriptional regulation, metabolites, and environmental signals [1,4]. Although this genome-based comparative analysis does not directly explain existing transcriptomic or phenotypic observations, it provides a genomic background for interpreting jellyfish metamorphosis. Several genes previously reported as candidates associated with metamorphosis, including *RXR* and *RDH-like* genes, were identified within the gene repertoires examined in this study, extending the range of retinoid-associated genes considered at the genomic level.

In this study, the identification of gene presence and copy number cannot directly inform functional activation or expression levels. In addition, assembly- and annotation-related biases may be present, and the availability or utilization of ligands for receptors such as RXR and AhR cannot be inferred from genomic data. Therefore, further studies integrating stage-specific transcriptomics, compound exposure experiments, and analyses of host-microbiome interactions will be required.

This study summarizes the current status of available Scyphozoa genome resources and presents the distribution of retinoid- and AhR-associated gene families. These results provide a reference framework for designing subsequent transcriptomic and experimental studies, rather than supporting direct functional conclusions.

## Author Contributions

Conceptualization: Y.-J.P., K.K.K.

Methodology: Y.-J.P., K.K.K.

Data Curation: Y.-J.P.

Formal Analysis: Y.-J.P.

Software: Y.-J.P.

Visualization: Y.-J.P.

Writing – Original Draft: Y.-J.P.

Writing – Review & Editing: Y.-J.P., K.K.K., Y.J.J., N.Y.L.

Supervision: S.S.Y., K.K.K.

Funding Acquisition: K.K.K.

## Data availability statement

All genome assemblies analyzed in this study are publicly available from previously published sources, and detailed information, including accession numbers, is provided in Table 1.

## Supporting Information captions

**Table S1**. Retinoid-related gene families identified in scyphozoan genomes.

**Table S2**. AhR-associated gene families identified in scyphozoan genomes.

**Table S3**. BUSCO completeness percentages for 21 scyphozoan genome assemblies.

